# Influence of different glycoproteins and of the virion core on SERINC5 antiviral activity

**DOI:** 10.1101/780577

**Authors:** William E. Diehl, Mehmet H. Guney, Pyae Phyo Kyawe, Judith M. White, Massimo Pizzato, Jeremy Luban

**Affiliations:** Program in Molecular Medicine, University of Massachusetts Medical School, 373 Plantation Street, Worcester MA 01605, USA; Department of Cell Biology, University of Virginia, Charlottesville, VA 22908, USA; Centre for Integrative Biology, University of Trento, 38123 Trento, Italy; Department of Biochemistry and Molecular Pharmacology, University of Massachusetts Medical School, Worcester, MA 01605, USA

## Abstract

Host plasma membrane protein SERINC5 is incorporated into budding retrovirus particles where it blocks subsequent entry into susceptible target cells. Three accessory proteins encoded by diverse retroviruses, HIV-1 Nef, EIAV S2, and MLV Glycogag, each independently disrupt SERINC5 antiviral activity, by redirecting SERINC5 from the site of virion assembly on the plasma membrane to an internal RAB7^+^ endosomal compartment. Pseudotyping retroviruses with particular glycoproteins, e.g., the vesicular stomatitis glycoprotein (VSV G), renders the infectivity of particles resistant to inhibition by virion-associated SERINC5. To better understand viral determinants for SERINC5-sensitivity, the effect of SERINC5 was assessed using HIV-1, MLV, and M-PMV virion cores, pseudotyped with glycoproteins from Arenavirus, Coronavirus, Filovirus, Rhabdovirus, Paramyxovirus, and Orthomyxovirus genera. Infectivity of particles, pseudotyped with HIV-1, amphotropic-MLV, or influenza virus glycoproteins, was decreased by SERINC5, whether the core was provided by HIV-1, MLV, or M-PMV. Particles generated by all three cores, and pseudotyped with glycoproteins from either avian leukosis virus-A, human endogenous retrovirus K (HERV-K), ecotropic-MLV, HTLV-1, Measles morbillivirus, lymphocytic choriomeningitis mammarenavirus (LCMV), Marburg virus, Ebola virus, severe acute respiratory syndrome-related coronavirus (SARS-CoV), or VSV, were insensitive to SERINC5. In contrast, particles pseudotyped with M-PMV, RD114, or rabies virus (RABV) glycoproteins were sensitive to SERINC5, but only with particular retroviral cores. Resistance to SERINC5 by particular glycoproteins did not correlate with reduced SERINC5 incorporation into particles or with the route of viral entry. These findings indicate that some non-retroviruses may be sensitive to SERINC5 and that, in addition to the viral glycoprotein, the retroviral core influences sensitivity to SERINC5.

**IMPORTANCE:** The importance of SERINC5 for inhibition of retroviruses is underscored by convergent evolution among three non-monophyletic retroviruses, each of which encodes a structurally unrelated SERINC5 inhibitor. One of these retroviruses causes tumors in mice, a second anemia in horses, and a third causes AIDS. SERINC5 is incorporated into retrovirus particles where it blocks entry into target cells, via a mechanism that is dependent on the viral glycoprotein. Here we demonstrate that retroviruses pseudotyped with glycoproteins from several non-retroviruses are also inhibited by SERINC5, suggesting that enveloped viruses other than retroviruses may also be inhibited by SERINC5. Additionally, we found that sensitivity to SERINC5 is determined by the retrovirus core, as well as by the glycoprotein. By better understanding how SERINC5 inhibits viruses we hope to extend fundamental understanding of virus replication and of the native role of SERINC5 in cells, and perhaps to advance the development of new antiviral strategies.

## INTRODUCTION

HIV-1 Nef is important for maximal HIV-1 replication *in vivo* and for progression to AIDS (1–3). Nef is a multifunctional accessory protein that downregulates CD4, MHC, and TCR from the cell surface (4–8). Nef also enhances HIV-1 infectivity in single-round infection experiments (9–16) by overcoming the antiviral effects of SERINC5 and SERINC3 (17, 18), though, of the two, SERINC5 is the more potent restriction factor. SERINC5 is incorporated into budding virions where it inhibits subsequent fusion of the virion membrane with target cell membranes. Nef counteracts SERINC5 by removing it from the cell surface so that it is not incorporated into nascent virions (17–20).

HIV-1 is not the only virus inhibited by SERINC5. SIVs lacking *nef* are also inhibited by SERINC5 and SIV *nef*s counteract this inhibition (17) with a potency that is proportional to the prevalence of SIV in wild primate populations (21). Two examples of convergent evolution of anti-SERINC function by virally encoded proteins are found outside of primate immunodeficiency viruses. Murine leukemia virus (MLV) Glycogag and equine infectious anemia virus (EIAV) S2 are viral antagonists of SERINC5 activity, and neither share sequence or structural homology with Nef, nor to each other (17, 22–24).

The mechanism by which virion-associated SERINC5 inhibits HIV-1 entry is unknown. The block is manifest after virion attachment to target cells, apparently at the stage of fusion pore expansion; virion contents mix with target cell cytoplasm but virion core transfer to the cytoplasm is inhibited (17, 19). Otherwise isogenic virions pseudotyped with HIV-1 Env glycoproteins from different HIV-1 isolates exhibit a range of dependency on Nef and of sensitivity to SERINC5 (25, 26). SERINC5 increases HIV-1 sensitivity to antibodies and peptides targeting the membrane-proximal external region of gp41, suggesting that it somehow alters the conformation of the HIV-1 glycoprotein near the virion membrane (19, 25). Importantly, HIV-1 particles pseudotyped with vesicular stomatitis virus (VSV) G or Ebola virus glycoprotein are resistant to SERINC5 antiviral activity (17, 18, 24). These initial observations suggest a correlation between the location of viral fusion and sensitivity to SERINC5 activity, with glycoproteins that mediate fusion at the cell surface (Env from HIV-1 and amphotropic MLV [A-MLV]) being sensitive and those that mediate fusion in endo-lysosomal compartments (VSV-G and Ebola GP) being resistant (17, 24). Taken together these results indicate that the virion glycoprotein is a viral determinant of sensitivity to SERINC5.

SERINC5 is a multipass transmembrane that localizes almost exclusively to the plasma membrane (17, 18). As such, in the absence of counter-measures, all enveloped viruses would be expected to encounter SERINC5 during viral egress, and to potentially be subject to its antiviral effects. We sought to address the breadth of SERINC5 antiviral activity and assess whether the route of entry impacts the sensitivity of viral glycoproteins to the antiviral effects of SERINC5. To do so, we investigated whether the co-expression of SERINC5 during viral production could inhibit a variety of glycoprotein pseudotypes of HIV, MLV, or M-PMV cores. Using this system, we tested the sensitivity of a number of retroviral Envs as well as representative glycoproteins from the Arenavirus, Coronavirus, Filovirus, Rhabdovirus, Paramyxovirus, and Orthomyxovirus genera. Consistent with previous findings, we observed that glycoprotein is a major determinant of SERINC5 sensitivity. While many glycoproteins were universally insensitive to the antiviral effects of SERINC5, the glycoproteins from NL4.3, A-MLV, and influenza were inhibited by SERINC5 in all viral core pseudotypes tested. No correlation was observed between SERINC5 sensitivity and the route of viral entry mediated by the viral glycoprotein. Unexpectedly, we also observed that sensitivity to SERINC5 antiviral activity for M-PMV, RD114, and rabies virus (RABV) glycoproteins depended on the retroviral core onto which they were pseudotyped. Our findings reveal that an interplay between virion core and glycoprotein determines the sensitivity to SERINC5 antiviral activity.

## RESULTS

To determine which viral glycoproteins are sensitive to the antiviral activity of SERINC5 we assessed infectivity of pseudotyped GFP-expressing lentiviral vectors produced in the presence or absence of SERINC5. Included in this panel was a diverse selection of retroviral Env glycoproteins, including those from human immunodeficiency virus-1 (HIV-1), avian leukosis virus A (ALV-A), human endogenous retrovirus K (HERV-K), feline endogenous retrovirus RD114, Mason-Pfizer monkey virus (M-PMV), ecotropic MLV (EcoMLV), amphotropic murine leukemia virus (A-MLV), and human T-cell lymphoma virus-1 (HTLV-1). We also tested the glycoproteins from an assortment of RNA viruses including influenza (H7/N1), parainfluenza 5 (PIV5), measles, rabies virus (RABV), lymphocytic choriomeningitis virus (LCMV), Marburg virus (MARV), Ebola virus Zaire [Mayinga] (EBOV), severe acute respiratory virus coronavirus (SARS CoV), and vesicular stomatitis virus (VSV). For these experiments we considered glycoprotein pseudotypes to be sensitive to SERINC5 restriction if viral titer was reduced at least 10-fold in the presence of SERINC5.

Similar to the findings of others (17, 18, 24), we observed that SERINC5 causes a greater than 100-fold reduction in viral infectivity for HIV-1 and A-MLV pseudotypes, while no significant reduction was observed for EBOV and VSV pseudotypes (Fig. 1A and Table 1). Interestingly, we observed >10-fold reduction of infectivity of H7/N1 influenza and RABV pseudotypes. No other pseudotypes displayed >10-fold reduced infectivity with SERINC5. These observations indicate that restriction by SERINC5 is not dictated by how the viral glycoprotein mediates fusion, as fusion mediated by influenza (27) or by RABV (28) occurs in a pH-dependent fashion in the endo-lysosomal compartment, while HIV-1-(29, 30), A-MLV-, M-PMV- and HTLV-1-mediated (31) fusion occurs in a pH-independent manner.

**Table 1.** Magnitude restriction of the indicated pseudotypes by SERINC5.

|  | <u>HIV Core</u> |  |  | <u>MLV Core</u> |  |  | <u>M-PMV Core</u> |  |  |
| --- | --- | --- | --- | --- | --- | --- | --- | --- | --- |
|  | Fold<br>Restriction | SEM | n | Fold<br>Restriction | SEM | n | Fold<br>Restriction | SEM | n |
| HIV-1 | **132.9** | **35.6** | **3** | **23.5** | **10.5** | **3** | **360.1** | **181.1** | **3** |
| ALV-A | 3.6 | 1.0 | 4 | 3.4 | 0.6 | 4 | 3.9 | 1.2 | 4 |
| HERV-K | 1.2 | 0.3 | 3 | 3.2 | 1.4 | 5 | 1.2 | 0.2 | 4 |
| RD114 | 4.6 | 0.7 | 4 | 7.4 | 2.5 | 5 | **24.8** | **10.9** | **6** |
| M-PMV | 1.5 | 0.2 | 3 | **46.3** | **18.6** | **6** | **104.2** | **31.8** | **8** |
| EcoMLV | 1.7 | 0.6 | 3 | 2.3 | 0.6 | 4 | 3.2 | 1.6 | 5 |
| A-MLV | **123.8** | **22.7** | **12** | **61.7** | **33.7** | **7** | **315.8** | **87.9** | **12** |
| HTLV-1 | 1.2 | 0.4 | 3 | 2.1 | 0.8 | 5 | 1.1 | 0.2 | 4 |
| Flu (H7) | **27.3** | **10.6** | **5** | **67.1** | **27.2** | **4** | **31.2** | **9.6** | **8** |
| PIV5 | 2.1 | 0.7 | 6 | 2.8 | 0.9 | 3 | 9.6 | 3.5 | 7 |
| Measles | 2.0 | 0.2 | 4 | 1.5 | 1 | 3 | 1 | 0.1 | 4 |
| RABV | **12.0** | **3.6** | **3** | **24.2** | **10.9** | **6** | 1.2 | 0.5 | 4 |
| LCMV | 4.1 | 1.1 | 3 | 1.2 | 0.1 | 3 | 5.1 | 2.4 | 7 |
| MARV | 1.9 | 0.5 | 3 | 2.6 | 0.4 | 3 | 3.2 | 0.8 | 4 |
| EBOV | 1.9 | 0.1 | 3 | 4.6 | 1.7 | 3 | 1.7 | 0.2 | 4 |
| SARS-CoV | 0.9 | 0.2 | 5 | 3.5 | 1.0 | 4 | 1 | 0.2 | 4 |
| VSV | 3.1 | 0.6 | 11 | 1.1 | 0.2 | 6 | 1.0 | 0.1 | 11 |

**Table 2.** List of expression plasmids used in this study.

| **Glycoprotein** | **Plasmid** | **Source** | **Addgene #** | **Reference** |
| --- | --- | --- | --- | --- |
| ALV EnvA | pCB6-EnvA | Judith White (Univ. of Virginia) | 74420 | (51) |
| LCMV Arm53b | pCMV WT GP | Juan de la Torre (Scripps Research) | N/A | (52) |
| C-term. truncated SARS CoV S | pKS-SARS-SΔ19 | Shuetsu Fukushi (National Institute of Infectious Diseases, Japan) | N/A | (53) |
| M-PMV Env | pTMT | Eric Hunter (Emory Univ.) | N/A | (54) |
| HTLV-1 Env | pV1/HTLV | Paul Bieniasz (Rockefeller Univ.) | N/A | (55) |
| MLV Eco Env | pHCMV-EcoEnv | Miguel Sena-Esteves (Univ. of Mass. Med. School) | 15802 | (56) |
| MLV Amphi Env | pHCMV-AmphiEnv | Miguel Sena-Esteves (Univ. of Mass. Med. School) | 15799 | (56) |
| HERV-K Env | pCAGGS-HERV-K | Sean Whelan (Harvard Univ.) | N/A | (57) |
| PIV5 F | pCAGGS-PIV5 F | Sean Whelan (Harvard Univ.) | N/A | (58) |
| PIV5 HN | pCAGGS-PIV5 HN | Sean Whelan (Harvard Univ.) | N/A | (58) |
| Flu HA<br>(H7/Kp<br>Rostock) | pFPV-HA | John Olsen (Univ. of North<br>Carolina Chapel Hill<br>[Emeritus]) | N/A | (59) |
| Flu NA<br>(H1N1;<br>A/PR/8/34) | pEF6-NA | John Olsen (Univ. of North<br>Carolina Chapel Hill<br>[Emeritus]) | N/A | (60) |
| Flu M2 | pCB6-M2 | John Olsen (Univ. of North<br>Carolina Chapel Hill<br>[Emeritus]) | N/A | (61) |
| C-term.<br>truncated<br>Measles F | pCG-FΔ30 | Els Verhoyen (Ecole<br>Normale Supérieure de<br>Lyon) | N/A | (62) |
| C-term.<br>truncated<br>Measles H | pCG-HΔ24 | Els Verhoyen (Ecole<br>Normale Supérieure de<br>Lyon) | N/A | (62) |
| RD114A Env<br>with Ampho<br>MLV Env<br>cytoplasmic<br>tail | phCMV-RD114/TR | François-Loïc Cosset (Ecole<br>Normale Supérieure de<br>Lyon) | N/A | (63) |
| VSV G | pMD2.G | Didier Trono (Ecole<br>Polytechnique Federale de<br>Lausanne) | 12259 |  |
| Rabies virus<br>G | pLTR-RVG | Jakob Reiser (US FDA) | 17577 | (64) |
| Mayinga<br>EBOV GP | pCB6-Zaire<br>EBOV | Graham Simmons (Univ. of<br>California, San Francisco) | N/A | (65) |
| Sudan Ebola<br>GP | pCB6-Sudan<br>EBOV | Graham Simmons (Univ. of<br>California, San Francisco) | N/A | (65) |
| Reston Ebola<br>GP | pCB6-Reston<br>EBOV | Graham Simmons (Univ. of<br>California, San Francisco) | N/A | (65) |
| Ravn MARV<br>GP | pCAGGS-MARV | Graham Simmons (Univ. of<br>California, San Francisco) | N/A | (66) |
| Bundibugyo<br>Ebola GP | pCAGGS-BDBV | Graham Simmons (Univ. of<br>California, San Francisco) | N/A | (67) |
| Tai Forest Ebola GP | pcDNA6-TAFV | Graham Simmons (Univ. of California, San Francisco) | N/A | (67) |
| Lloviu Ebola GP | pCAGGS-LLOV | Graham Simmons (Univ. of California, San Francisco) | N/A | (67) |
| Makona EBOV GP | pGL4.23 WT 2014 EBOV ΔMuc | N/A | 86021 | (36) |
| A82V Makona EBOV GP | pGL4.23 A82V 2014 EBOV ΔMuc | N/A | 86022 | (36) |
| C-term. truncated NL4.3 Env (C/O) | pcDNA-NL4.3 Env ΔCT | N/A | Pending | This Work |
| GPI anchored TVA | pCMMP-TVA800 | Edward Callaway (Salk Institute) | 15778 | (68) |
| murine CAT1 | pBABE-puro-mCAT1 | Massimo Pizzato (Univ. of Trento) | N/A | (23) |

**Figure 1.**
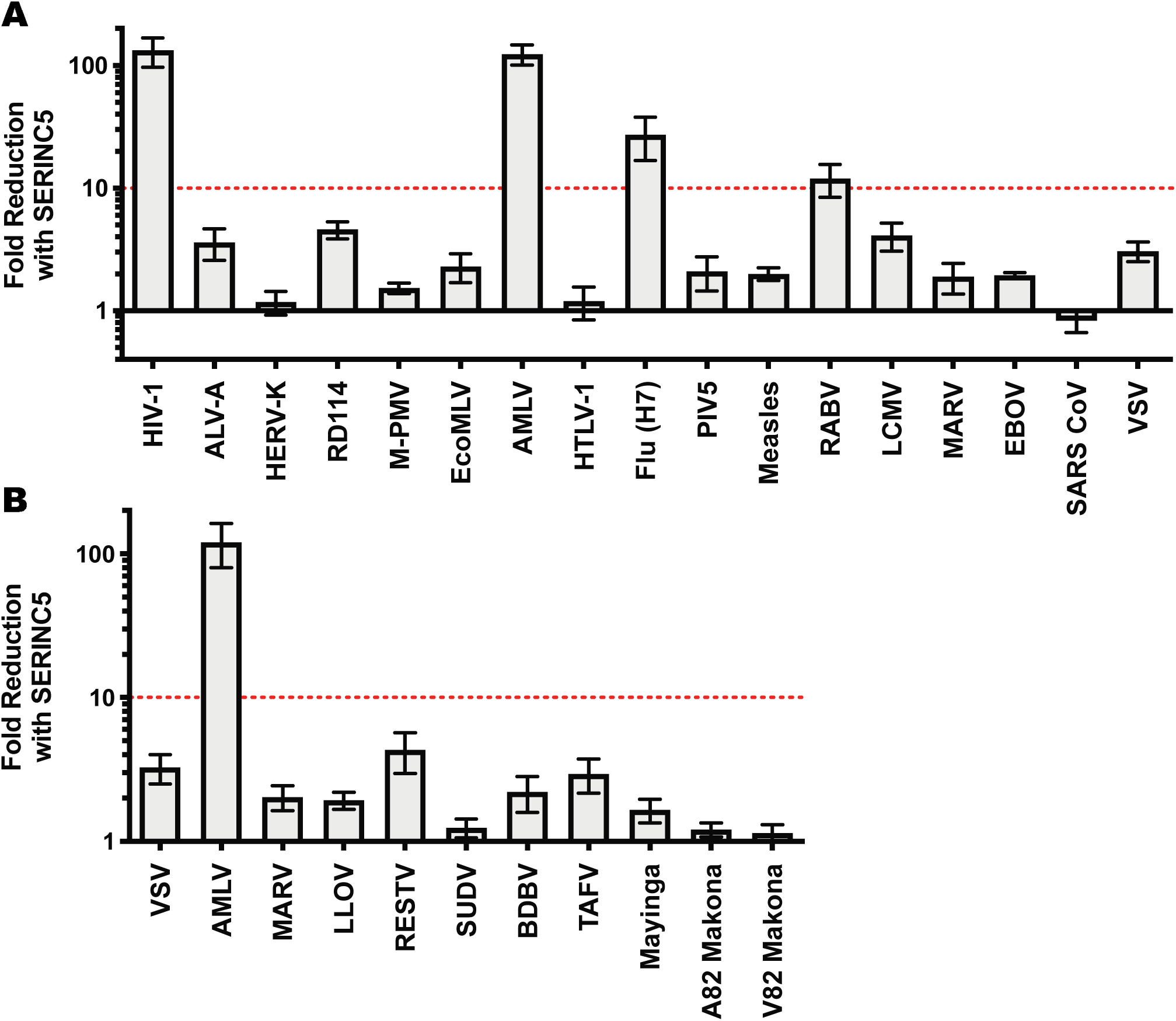
Sensitivity of HIV-1 pseudotypes to SERINC5 antiviral activity. (A) Effect of SERINC5 on transduction efficiency by HIV-1 cores pseudotyped with a diverse panel of viral glycoproteins. With the exception of HIV-1 pseudotypes, transductions were conducted in HEK293 cells. For transductions involving ecotropic MLV or avian leukosis virus A pseudotypes, target HEK293 cells were transfected with viral receptor prior to transductions. TZMbl cells were used to evaluate infectivity of HIV-1 pseudotypes. (B) Sensitivity of filoviral glycoprotein pseudoviruses to SERINC5. Plotted is the difference in infectivity between virus produced in the absence versus the presence of SERINC5. Each condition shows results of vector production from at least three independent transfections. The red lines indicates 10-fold lower infectivity in the presence of SERINC5, our arbitrary cut off for SERINC sensitivity. HIV-1: human immunodeficiency virus-1, ALV-A: avian leukosis virus A, HERV-K: human endogenous retrovirus K, RD114: feline endogenous retrovirus RD114, M-PMV: Mason-Pfizer monkey virus, EcoMLV: ecotropic MLV, AMLV: amphotropic MLV, HTLV-1: human T-cell lymphotropic virus-1, Flu: influenza type A, PIV5: parainfluenza virus 5, RABV: rabies virus, LCMV: Lymphocytic choriomeningitis virus, MARV: Marburg virus, EBOV: Mayinga isolate of Zaire ebolavirus, SARS CoV: severe acute respiratory syndrome coronavirus, VSV: vesicular stomatitis virus, LLOV: Lloviu virus, RESTV: Reston virus, SUDV: Sudan virus, BDBV: Bundibugyo virus, TAFV: Taï Forest virus, Mayinga: Mayinga isolate of Ebola virus, Makona: Makona isolate of Ebola virus.

Next we tested a panel of filoviral glycoproteins for sensitivity to SERINC5 restriction. All of these glycoproteins require proteolytic processing (32, 33) following internalization into the target cell and utilize the lysosomal protein NPC1 to initiate viral fusion (34, 35). In addition to the EBOV and MARV glycoproteins tested in Fig 1A, this panel included glycoproteins from Bundibugyo (BDBV), Lloviu (LLOV), Reston (RESTV), Sudan (SUDV), Taï Forest (TAFV), and the 2014 Makona glycoprotein variant (A82) that initiated the 2013-2016 outbreak, along with an infectivity-enhancing derivative (GP-A82V) that arose during the outbreak (36, 37). As shown in Fig. 1B, none of the filoviral glycoproteins were inhibited >10-fold in the presence of SERINC5. However, there may be modest differences in sensitivity to SERINC5 activity, specifically RESTV and TAFV GP appear slightly more sensitive (4.3- and 2.9-fold, respectively) to SERINC5 inhibition compared to either Mayinga or Makona Ebola virus glycoproteins (1.65- and 1.2-fold, respectively).

HIV-1 Nef, MLV glygoGag, and EIAV S2 counteract SERINC5 antiviral activity by removing SERINC5 protein from the cell surface and relocalizing it to an endosomal compartment (17, 18, 22). The ability of a viral glycoprotein to re-localize a normally plasma membrane localized antiviral protein has been previously shown for HIV-2 Env and human BST2 (38). Thus, we reasoned that viral glycoproteins may confer resistance to SERINC5 activity by re-localizing SERINC5 to an internal membrane compartment. To test this, we compared SERINC5 incorporation into HIV-1 virus-like particles (VLPs) pseudotyped with the various glycoproteins shown in Fig 1A. We found that HIV-1 VLPs universally incorporated SERINC5 irrespective of the viral glycoprotein present (Figure 2). In replicate blotting, only HERV-K Env showed a consistently lower level of SERINC5 incorporation into viral particles (data not shown). However, this observation is likely to be caused by pleiotropic effects of cells transfected with this glycoprotein, as cell growth was significantly reduced compared to other transfections, and reduced levels of Gag and GFP were also observed (Fig. 2, data not shown). Regardless, no direct correlation between SERINC5 exclusion from virions and resistance to its antiviral effects was evident.

**Figure 2.**
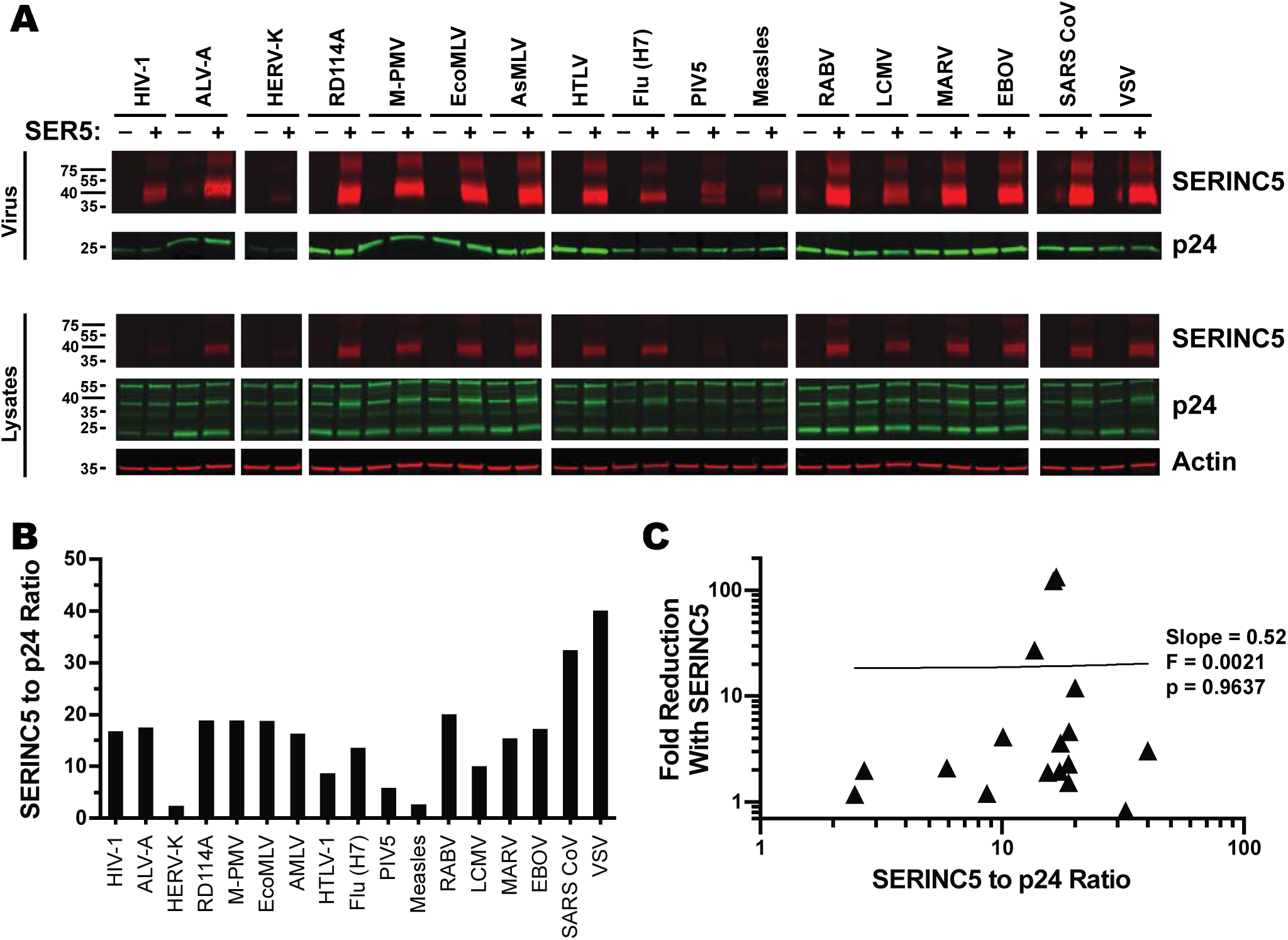
Incorporation of SERINC5 into HIV-1 pseudovirus does not correlate with sensitivity to its antiviral effects. (Top) Western blots of purified HIV-1 pseudovirions produced in the presence or absence of C-terminally HA-tagged SERINC5. Blots were probed with mouse monoclonal anti-HA and human anti-p24 monoclonal 241-D. (Bottom) Western blots of lysates from producer HEK293s of the pseudovirions shown above. Blots were probed with mouse anti-actin in addition to anti-HA and anti-p24 used for purified pseudovirions.

A previous report indicated that MLV virions pseudotyped with RD114 Env are susceptible to the antiviral effects of SERINC5 (24). However, in the presence of SERINC5 we only observed a modest ∼4.5-fold reduction in viral titer of RD114 pseudotyped HIV-1 virions (Fig. 1A). In response to this discrepancy, we sought to determine if the viral core modulates susceptibility to SERINC5 antiviral activity. Thus, we tested the same panel of glycoproteins for SERINC5 sensitivity when pseudotyped on different virion cores. First, we tested the SERINC5 sensitivity of the same panel of glycoproteins as in Fig. 1A on MLV viral cores (Figure 3A and Table 1). We observed that the glycoproteins sensitive to SERINC5 restriction on HIV-1 cores (HIV-1, A-MLV, Flu, and Rabies) were also restricted when pseudotyped on MLV cores. Additionally, we observed that M-PMV Env was sensitive to SERINC restriction when pseudotyped onto MLV cores, whereas it was not when pseudotyped onto HIV-1 cores. Returning to the initial impetus for exploring different cores, we observed a ∼7.5-fold reduction of infectivity for RD114 pseudotyped MLV cores when produced in the presence of SERINC5, which is similar to the magnitude of the inhibitory effect reported by Ahi et al. (24).

**Figure 3.**
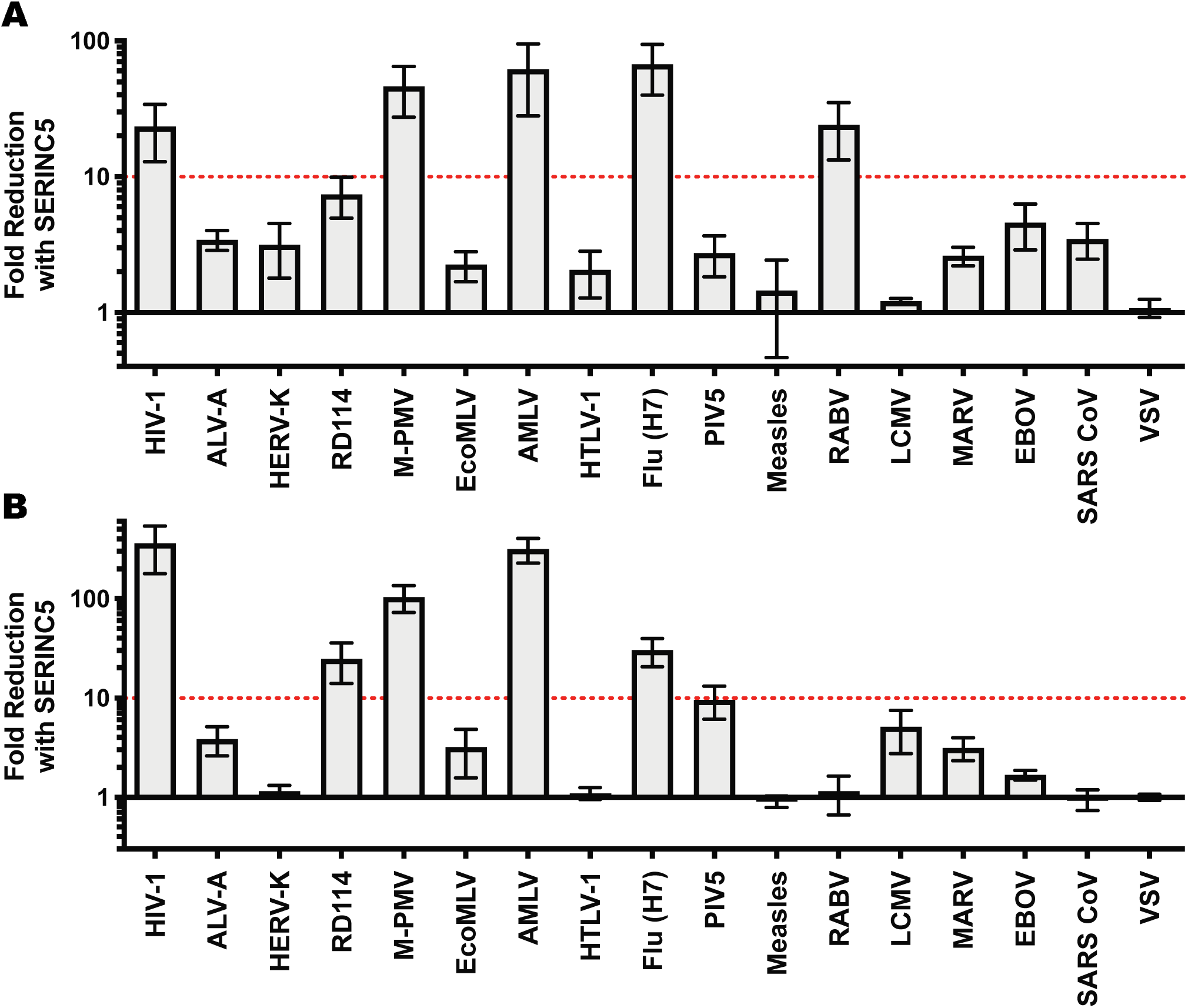
Differential sensitivity of glycoproteins to SERINC5 antiviral activity based on the vial core onto which they are pseudotyped. Effects of SERINC5 on the HEK293 infectivity of a diverse panel of viral glycoproteins pseudotyped on (A) MLV or (B) M-PMV cores. Infectivity was assessed as described in Fig. 1. Plotted is the difference in infectivity between virus produced in the absence versus the presence of SERINC5 from at least three independent transfections. The red line indicates 10-fold lower infectivity in the presence of SERINC5, our arbitrary cut off for SERINC sensitivity.

Due to observed differences in SERINC5 sensitivity with M-PMV pseudotypes of HIV-1 and MLV cores, we next tested for SERINC5 antiviral activity against our panel of glycoproteins pseudotyped onto M-PMV cores (Figure 3B and Table 1). These experiments showed that the infectivity of M-PMV cores bearing M-PMV Env was significantly reduced in the presence of SERINC5. Furthermore, we observed that pseudotypes of M-PMV cores with HIV-1, A-MLV, and Flu glycoproteins were sensitive to SERINC5 restriction, similar to the observations with both HIV-1 and MLV cores. In contrast to observations with HIV-1 and MLV cores, the SERINC5-mediated reduction of infectivity for M-PMV cores pseudotyped with RD114 (∼20-fold) surpassed our 10-fold cutoff for significance. Conversely, rabies virus (RABV) glycoprotein pseudotypes of M-PMV were unaffected by the antiviral effects of SERINC5, in contrast to what was observed for RABV glycoprotein pseudotyped HIV-1 and MLV cores. Finally, SERINC5 reduced the infectivity of PIV5 pseudotyped M-PMV cores by ∼9.5-fold, just under our 10-fold significance cutoff.

## DISCUSSION

Initial reports indicated that the viral glycoprotein is a determinant of sensitivity to SERINC5 antiviral activity (17, 18, 25, 26) and suggested that viral glycoproteins which mediate fusion via a pH-dependent, endocytic entry pathway are resistant to SERINC5 antiviral activity (17, 24). Here, to examine these issues further, pseudotypes using glycoproteins from diverse families of enveloped viruses were assessed for sensitivity to restriction by SERINC5. We observed that SERINC5 restricted virions pseudotyped with glycoproteins from several retroviruses (HIV-1, A-MLV, RD114, and M-PMV), influenza A (Orthomyxoviridae), and rabies (Rhabdoviridae). To our knowledge, this is the first time antiviral activity of SERINC5 has been described for a non-retroviral glycoprotein. As the glycoproteins of these viruses were studied as retroviral pseudotypes, it remains to be established if the infectivity of authentic influenza or rabies viruses are affected by SERINC5, or other SERINC family members. Additionally, our observation with influenza A and rabies glycoproteins demonstrates that mediating entry via an endocytic route does not, in itself, protect from the antiviral effects of SERINC5.

While Env glycoproteins from the retroviruses HIV-1, MLV, and RD114 have all previously been found to be inhibited by SERINC5 (17, 18), we now report that M-PMV glycoprotein is SERINC5-sensitive as well. Interestingly, we saw a ∼100-fold reduction in infectivity of autologously pseudotyped M-PMV cores when produced in the presence of SERINC5. This observation was unexpected given that lenti- and gammaretroviruses encode accessory factors that counteract SERINC5 activity. And yet, functionally intact M-PMV (the only viral gene known to be missing from the GFP-expressing M-PMV vector is Env, which is complemented *in trans* during the transfection) was sensitive to the antiviral effects of human SERINC5.

Surprisingly, we observed that particular glycoproteins displayed different sensitivity to the antiviral effects of SERINC5 depending on the viral core onto which they were pseudotyped. For instance, rabies virus glycoprotein was inhibited by SERINC5 when on HIV-1 or MLV cores, but insensitive to SERINC5 when on M-PMV cores. In contrast, M-PMV glycoprotein was sensitive to SERINC5 restriction when on MLV or M-PMV cores, but resistant when on HIV-1 cores. Additionally, a ∼17-fold SERINC5-mediated inhibition was observed for M-PMV cores pseudotyped with RD114 Env, while 7.5-fold and 4.3-fold inhibitions were observed for RD114 pseudotypes of MLV and HIV cores, respectively. A previous report demonstrated similar magnitude inhibition for RD114-pseudotyped MLV by endogenous SERINC activity (24). Regardless, RD114 showed little sensitivity to SERINC5 when pseudotyped onto HIV-1 cores. Neutralization by monoclonal antibodies that target the membrane-proximal domain of HIV-1 glycoprotein is altered by the presence of SERINC5 (19, 25). Given that MA, the membrane proximal domain of *gag* makes contacts with the retroviral TM (40, 41), one can imagine that SERINC5 has the potential to influence interactions between MA and TM in the HIV-1 virion that are essential for infectivity. In similar fashion, SERINC5 might influence retroviral core interactions by the heterologous glycoproteins tested here, for which SERINC5 restriction activity was core-dependent, i.e., the rabies virus, MPMV, and RD114 glycoproteins.

## MATERIALS AND METHODS

### Plasmid DNA

Plasmids used in this study are described in Table 1, including Addgene or NIH AIDS Reagent Program code numbers (where applicable), where full plasmid sequences can be obtained. A pcDNA3.1 based vector bearing codon-optimized pNL4-3 *env* with a cytoplasmic tail truncation after residue 710 (HXB2 residue 712), similar to that previously described (42), was generated using standard cloning techniques and is available from Addgene.

### Cell culture

HEK293 cells were obtained from the ATCC. The HIV indicator cell line TZM-bl (Cat#8129) was obtained from the AIDS Research and Reference Reagent Program (Division of AIDS, NIAID, NIH) and were deposited by Drs. John C. Kappes and Xiaoyun Wu (43). Both cell lines were maintained in DMEM supplemented with 10% FBS and 10 mM HEPES.

### Virus production, and transductions

All viral stocks were generated by Mirus TransIT-LT1 (Mirus Bio, Madison, WI) mediated transfection of HEK293 cells. 12 well plates were seeded with 3×10^5^ cells per well 24 hours prior to transfection. In the evening of the following day, 3.375 μl LT1 reagent was used to transfect plasmids as follows: For the production of pseudotyped HIV-1 virions 625 ng pNL-EGFP/CMV⍰WPREΔU3 (44) and 465 ng pCD/NL-BH*ΔΔΔ (45) were co-transfected with 155 ng glycoprotein expression vector. For the production of pseudotyped MLV virions 625 ng pLXIN-GFP (46) and 465 ng pCS2+mGP (47) were co-transfected with 155 ng glycoprotein expression vector. For the production of pseudotyped M-PMV virions, 1090 ng pSARM-EGFP (48) was co-transfected with 155 ng glycoprotein expression vector. In all cases, either 100ng pcDNA-SERINC5 (17) or 110 ng empty pcDNA3.1 (Thermo Fisher Scientific, Waltham, MA) vector was included in these transfections. The morning following transfection medium was replaced with fresh DMEM and virus containing supernatant was harvested 48 hours after media change. This supernatant was spun for 10 minutes at 2,500 x *g* to remove cellular debris and stored at 4°C until used for transduction.

HEK293 or TZMbl cells were seeded at 1×10^5^ or 5×10^5^, respectively, in 12-well plates 24 hours prior to transduction. For experiments involving ecotropic MLV or avian leukosis virus A, HEK293 cells were transfected in 6-well plates with 2.5 μg of pBABE-puro-mCAT or pCMMP-TVA800 using TransIT-LT1 and the subsequent day these transfected cells were split and plated for transductions. For transductions, culture supernatant was replaced with three dilutions of virus containing supernatant and incubated overnight at 37°C. Virus containing medium was replaced and cells were incubated for an additional 48 hours, following which they were trypsinized and assessed for GFP expression via fluorescent activated cell sorting using the Accuri C6 (BD Biosciences, San Jose, CA). Analysis was performed using FlowJo Macintosh v10.1 (FlowJo, LLC, Ashland, OR).

### Virion purification and western blotting

Viral pseudotypes were produced as above, except transfections were performed in 6-well plates so the number of cells plated and DNA introduced were doubled. The resulting virus-containing supernatant was overlayed on 20% sucrose in TNE buffer (50 mM TRIS, 100 mM NaCl, 0.1 mM EDTA, pH7.4) and viruses were pelleted via ultracentrifugation for 2 hours at 125,000 x *g* at 4°C using an SW55-Ti rotor (Beckman Coulter, Indianapolis, IN). Following centrifugation, tubes were washed with 1 ml of ice cold PBS and viral pellets were directly lysed in 50 μl 2x Laemmli buffer containing 50 mM TCEP [Tris(2-carboxyethyl)phosphine] incubated at room temp for 5 minutes. Cell lysates were prepared in parallel by washing transfected HEK293s once with 1 ml ice cold PBS, detaching from the plate by scraping, pelleting, and subsequently lysing for 20 minutes on ice in 150 μl SERINC lysis buffer (10 mM HEPES, pH 7.5, 100 mM NaCl, 1 mM TCEP [Tris(2-carboxyethyl)phosphine], 1% DDM [n-Dodecyl-β-D-maltoside]) containing cOmplete mini protease inhibitor (Sigma-Aldrich, St. Louis, MO). Lysates were clarified by centrifugation for 5 minutes at 10,000 x *g* and 4°C, following which supernatants were transferred to a new centrifuge tube and protein content was quantified via Reducing Agent Compatible BCA Assay (Thermo Scientific, Waltham, MA) Volumes of lysate corresponding to equal protein content were combined 1:1 with 2x Laemmli buffer containing 50 mM TCEP and incubated at room temp for 5 minutes.

One half of the denatured viral pellet and approximately 8 μg protein from cellular lysates were run on 4-15% gradient acrylamide gels, and transferred to nitrocellulose membranes. SERINC5 levels were assessed via C-terminal HA tag using the mouse monoclonal HA.11 (Biolegend, San Diego, CA) at 1 μg/ml in Odyssey blocking buffer (LI-COR Biotechnology, Lincoln, NE). HIV-1 p24 was detected using human monoclonal antibody 241-D (49) at a concentration of 1 μg/ml in Odyssey blocking buffer. MLV p30 was detected with rat monoclonal antibody R187 (50) from unpurified culture medium following five days of culturing the R187 hybridoma (ATCC, Manassas, VA). This medium was diluted 1:200 in Odyssey blocking buffer. Cellular actin was detected using mouse anti-actin monoclonal ACTN05 (C4) (Abcam, Cambridge, MA) at a concentration of 0.5 μg/ml in Odyssey blocking buffer. All blots were developed using 1:10,000 dilutions of 680RD or 800CW fluorescently tagged secondary antibodies (LI-COR Biotechnology, Lincoln, NE) in Odyssey blocking buffer. Imaging of blots was performed using an Odyssey CLx system (LI-Cor Biotechnology) at a resolution of 84 μm using the ‘high quality’ setting. Quantitation of bands was done using the box tool in the Odyssey software package with adjacent pixels to the box serving as reference background levels for background subtraction.

## ACKNOWLEDGEMENTS

The following reagent was obtained through the NIH AIDS Reagent Program, Division of AIDS, NIAID, NIH: Anti-HIV-1 p24 Monoclonal (241-D) from Dr. Susan Zolla-Pazner. This work was supported by NIH grants DP1DA034990, RO1AI117839, and 1R37AI147868 to J.L.

## REFERENCES

1. Deacon NJ, Tsykin A, Solomon A, Smith K, Ludford-Menting M, Hooker DJ, McPhee DA, Greenway AL, Ellett A, Chatfield C, Lawson VA, Crowe S, Maerz A, Sonza S, Learmont J, Sullivan JS, Cunningham A, Dwyer D, Dowton D, Mills J. 1995. Genomic structure of an attenuated quasi species of HIV-1 from a blood transfusion donor and recipients. Science 270:988–991.

2. Kestler HW 3rd, Ringler DJ, Mori K, Panicali DL, Sehgal PK, Daniel MD, Desrosiers RC. 1991. Importance of the nef gene for maintenance of high virus loads and for development of AIDS. Cell 65:651–662.

3. Kirchhoff F, Greenough TC, Brettler DB, Sullivan JL, Desrosiers RC. 1995. Absence of Intact nef Sequences in a Long-Term Survivor with Nonprogressive HIV-1 Infection. N Engl J Med 332:228–232.

4. Anderson S, Shugars DC, Swanstrom R, Garcia JV. 1993. Nef from primary isolates of human immunodeficiency virus type 1 suppresses surface CD4 expression in human and mouse T cells. J Virol 67:4923–4931.

5. Mariani R, Skowronski J. 1993. CD4 down-regulation by nef alleles isolated from human immunodeficiency virus type 1-infected individuals. Proc Natl Acad Sci U S A 90:5549–5553.

6. Garcia JV, Miller AD. 1991. Serine phosphorylation-independent downregulation of cell-surface CD4 by nef. Nature 350:508–511.

7. Stove V, Van de Walle I, Naessens E, Coene E, Stove C, Plum J, Verhasselt B. 2005. Human immunodeficiency virus Nef induces rapid internalization of the T-cell coreceptor CD8alphabeta. J Virol 79:11422–11433.

8. Schwartz O, Maréchal V, Le Gall S, Lemonnier F, Heard JM. 1996. Endocytosis of major histocompatibility complex class I molecules is induced by the HIV-1 Nef protein. Nat Med 2:338–342.

9. Chowers MY, Spina CA, Kwoh TJ, Fitch NJ, Richman DD, Guatelli JC. 1994. Optimal infectivity in vitro of human immunodeficiency virus type 1 requires an intact nef gene. J Virol 68:2906–2914.

10. Schaeffer E, Geleziunas R, Greene WC. 2001. Human immunodeficiency virus type 1 Nef functions at the level of virus entry by enhancing cytoplasmic delivery of virions. J Virol 75:2993–3000.

11. Tobiume M, Lineberger JE, Lundquist CA, Miller MD, Aiken C. 2003. Nef Does Not Affect the Efficiency of Human Immunodeficiency Virus Type 1 Fusion with Target Cells. J Virol 77:10645–10650.

12. Campbell EM, Nunez R, Hope TJ. 2004. Disruption of the actin cytoskeleton can complement the ability of Nef to enhance human immunodeficiency virus type 1 infectivity. J Virol 78:5745–5755.

13. Cavrois M, Neidleman J, Yonemoto W, Fenard D, Greene WC. 2004. HIV-1 virion fusion assay: uncoating not required and no effect of Nef on fusion. Virology 328:36–44.

14. Day JR, Münk C, Guatelli JC. 2004. The membrane-proximal tyrosine-based sorting signal of human immunodeficiency virus type 1 gp41 is required for optimal viral infectivity. J Virol 78:1069–1079.

15. Schwartz O, Maréchal V, Danos O, Heard JM. 1995. Human immunodeficiency virus type 1 Nef increases the efficiency of reverse transcription in the infected cell. J Virol 69:4053–4059.

16. Aiken C, Trono D. 1995. Nef stimulates human immunodeficiency virus type 1 proviral DNA synthesis. J Virol 69:5048–5056.

17. Rosa A, Chande A, Ziglio S, De Sanctis V, Bertorelli R, Goh SL, McCauley SM, Nowosielska A, Antonarakis SE, Luban J, Santoni FA, Pizzato M. 2015. HIV-1 Nef promotes infection by excluding SERINC5 from virion incorporation. Nature 526:212–217.

18. Usami Y, Wu Y, Göttlinger HG. 2015. SERINC3 and SERINC5 restrict HIV-1 infectivity and are counteracted by Nef. Nature 526:218–223.

19. Sood C, Marin M, Chande A, Pizzato M, Melikyan GB. 2017. SERINC5 protein inhibits HIV-1 fusion pore formation by promoting functional inactivation of envelope glycoproteins. J Biol Chem 292:6014–6026.

20. Trautz B, Pierini V, Wombacher R, Stolp B, Chase AJ, Pizzato M, Fackler OT. 2016. The Antagonism of HIV-1 Nef to SERINC5 Particle Infectivity Restriction Involves the Counteraction of Virion-Associated Pools of the Restriction Factor. J Virol 90:10915–10927.

21. Heigele A, Kmiec D, Regensburger K, Langer S, Peiffer L, Stürzel CM, Sauter D, Peeters M, Pizzato M, Learn GH, Hahn BH, Kirchhoff F. 2016. The Potency of Nef-Mediated SERINC5 Antagonism Correlates with the Prevalence of Primate Lentiviruses in the Wild. Cell Host Microbe 20:381–391.

22. Chande A, Cuccurullo EC, Rosa A, Ziglio S, Carpenter S, Pizzato M. 2016. S2 from equine infectious anemia virus is an infectivity factor which counteracts the retroviral inhibitors SERINC5 and SERINC3. Proc Natl Acad Sci U S A 113:13197–13202.

23. Pizzato M. 2010. MLV glycosylated-Gag is an infectivity factor that rescues Nef-deficient HIV-1. Proc Natl Acad Sci U S A 107:9364–9369.

24. Ahi YS, Zhang S, Thappeta Y, Denman A, Feizpour A, Gummuluru S, Reinhard B, Muriaux D, Fivash MJ, Rein A. 2016. Functional Interplay Between Murine Leukemia Virus Glycogag, Serinc5, and Surface Glycoprotein Governs Virus Entry, with Opposite Effects on Gammaretroviral and Ebolavirus Glycoproteins. MBio 7:e01985–16.

25. Lai RPJ, Yan J, Heeney J, McClure MO, Göttlinger H, Luban J, Pizzato M. 2011. Nef decreases HIV-1 sensitivity to neutralizing antibodies that target the membrane-proximal external region of TMgp41. PLoS Pathog 7:e1002442.

26. Usami Y, Göttlinger H. 2013. HIV-1 Nef responsiveness is determined by Env variable regions involved in trimer association and correlates with neutralization sensitivity. Cell Rep 5:802–812.

27. Doms RW, Gething MJ, Henneberry J, White J, Helenius A. 1986. Variant influenza virus hemagglutinin that induces fusion at elevated pH. J Virol 57:603–613.

28. Piccinotti S, Kirchhausen T, Whelan SPJ. 2013. Uptake of rabies virus into epithelial cells by clathrin-mediated endocytosis depends upon actin. J Virol 87:11637–11647.

29. Stein BS, Gowda SD, Lifson JD, Penhallow RC, Bensch KG, Engleman EG. 1987. pH-independent HIV entry into CD4-positive T cells via virus envelope fusion to the plasma membrane. Cell 49:659–668.

30. McClure MO, Marsh M, Weiss RA. 1988. Human immunodeficiency virus infection of CD4-bearing cells occurs by a pH-independent mechanism. EMBO J 7:513–518.

31. McClure MO, Sommerfelt MA, Marsh M, Weiss RA. 1990. The pH independence of mammalian retrovirus infection. J Gen Virol 71 (Pt 4):767–773.

32. Chandran K, Sullivan NJ, Felbor U, Whelan SP, Cunningham JM. 2005. Endosomal proteolysis of the Ebola virus glycoprotein is necessary for infection. Science 308:1643–1645.

33. Gnirss K, Kühl A, Karsten C, Glowacka I, Bertram S, Kaup F, Hofmann H, Pöhlmann S. 2012. Cathepsins B and L activate Ebola but not Marburg virus glycoproteins for efficient entry into cell lines and macrophages independent of TMPRSS2 expression. Virology 424:3–10.

34. Carette JE, Raaben M, Wong AC, Herbert AS, Obernosterer G, Mulherkar N, Kuehne AI, Kranzusch PJ, Griffin AM, Ruthel G, Dal Cin P, Dye JM, Whelan SP, Chandran K, Brummelkamp TR. 2011. Ebola virus entry requires the cholesterol transporter Niemann-Pick C1. Nature 477:340–343.

35. Côté M, Misasi J, Ren T, Bruchez A, Lee K, Filone CM, Hensley L, Li Q, Ory D, Chandran K, Cunningham J. 2011. Small molecule inhibitors reveal Niemann-Pick C1 is essential for Ebola virus infection. Nature 477:344–348.

36. Diehl WE, Lin AE, Grubaugh ND, Carvalho LM, Kim K, Kyawe PP, McCauley SM, Donnard E, Kucukural A, McDonel P, Schaffner SF, Garber M, Rambaut A, Andersen KG, Sabeti PC, Luban J. 2016. Ebola Virus Glycoprotein with Increased Infectivity Dominated the 2013-2016 Epidemic. Cell 167:1088–1098.e6.

37. Urbanowicz RA, McClure CP, Sakuntabhai A, Sall AA, Kobinger G, Müller MA, Holmes EC, Rey FA, Simon-Loriere E, Ball JK. 2016. Human Adaptation of Ebola Virus during the West African Outbreak. Cell 167:1079–1087.e5.

38. Le Tortorec A, Neil SJD. 2009. Antagonism to and intracellular sequestration of human tetherin by the human immunodeficiency virus type 2 envelope glycoprotein. J Virol 83:11966–11978.

39. Li M, Waheed AA, Yu J, Zeng C, Chen H-Y, Zheng Y-M, Feizpour A, Reinhard BM, Gummuluru S, Lin S, Freed EO, Liu S-L. 2019. TIM-mediated inhibition of HIV-1 release is antagonized by Nef but potentiated by SERINC proteins. Proc Natl Acad Sci U S A.

40. Freed EO, Martin MA. 1996. Domains of the human immunodeficiency virus type 1 matrix and gp41 cytoplasmic tail required for envelope incorporation into virions. J Virol 70:341–351.

41. West JT, Weldon SK, Wyss S, Lin X, Yu Q, Thali M, Hunter E. 2002. Mutation of the dominant endocytosis motif in human immunodeficiency virus type 1 gp41 can complement matrix mutations without increasing Env incorporation. J Virol 76:3338–3349.

42. Wilk T, Pfeiffer T, Bosch V. 1992. Retained in vitro infectivity and cytopathogenicity of HIV-1 despite truncation of the C-terminal tail of the env gene product. Virology 189:167–177.

43. Wei X, Decker JM, Liu H, Zhang Z, Arani RB, Kilby JM, Saag MS, Wu X, Shaw GM, Kappes JC. 2002. Emergence of resistant human immunodeficiency virus type 1 in patients receiving fusion inhibitor (T-20) monotherapy. Antimicrob Agents Chemother 46:1896–1905.

44. Diehl WE, Stansell E, Kaiser SM, Emerman M, Hunter E. 2008. Identification of postentry restrictions to Mason-Pfizer monkey virus infection in New World monkey cells. J Virol 82:11140–11151.

45. Zhang X-Y, La Russa VF, Bao L, Kolls J, Schwarzenberger P, Reiser J. 2002. Lentiviral vectors for sustained transgene expression in human bone marrow-derived stromal cells. Mol Ther 5:555–565.

46. Newman RM, Hall L, Connole M, Chen G-L, Sato S, Yuste E, Diehl W, Hunter E, Kaur A, Miller GM, Johnson WE. 2006. Balancing selection and the evolution of functional polymorphism in Old World monkey TRIM5alpha. Proc Natl Acad Sci U S A 103:19134–19139.

47. Yamashita M, Emerman M. 2004. Capsid is a dominant determinant of retrovirus infectivity in nondividing cells. J Virol 78:5670–5678.

48. Diehl WE, Stansell E, Kaiser SM, Emerman M, Hunter E. 2008. Identification of postentry restrictions to Mason-Pfizer monkey virus infection in New World monkey cells. J Virol 82:11140–11151.

49. Gorny MK, Gianakakos V, Sharpe S, Zolla-Pazner S. 1989. Generation of human monoclonal antibodies to human immunodeficiency virus. Proc Natl Acad Sci U S A 86:1624–1628.

50. Chesebro B, Britt W, Evans L, Wehrly K, Nishio J, Cloyd M. 1983. Characterization of monoclonal antibodies reactive with murine leukemia viruses: use in analysis of strains of friend MCF and Friend ecotropic murine leukemia virus. Virology 127:134–148.

51. Gilbert JM, Bates P, Varmus HE, White JM. 1994. The receptor for the subgroup A avian leukosis-sarcoma viruses binds to subgroup A but not to subgroup C envelope glycoprotein. J Virol 68:5623–5628.

52. Saunders AA, Ting JPC, Meisner J, Neuman BW, Perez M, de la Torre JC, Buchmeier MJ. 2007. Mapping the landscape of the lymphocytic choriomeningitis virus stable signal peptide reveals novel functional domains. J Virol 81:5649–5657.

53. Fukushi S, Mizutani T, Saijo M, Matsuyama S, Miyajima N, Taguchi F, Itamura S, Kurane I, Morikawa S. 2005. Vesicular stomatitis virus pseudotyped with severe acute respiratory syndrome coronavirus spike protein. J Gen Virol 86:2269–2274.

54. Song C, Dubay SR, Hunter E. 2003. A tyrosine motif in the cytoplasmic domain of mason-pfizer monkey virus is essential for the incorporation of glycoprotein into virions. J Virol 77:5192–5200.

55. Cowan S, Hatziioannou T, Cunningham T, Muesing MA, Gottlinger HG, Bieniasz PD. 2002. Cellular inhibitors with Fv1-like activity restrict human and simian immunodeficiency virus tropism. Proc Natl Acad Sci U S A 99:11914–11919.

56. Sena-Esteves M, Tebbets JC, Steffens S, Crombleholme T, Flake AW. 2004. Optimized large-scale production of high titer lentivirus vector pseudotypes. J Virol Methods 122:131–139.

57. Robinson LR, Whelan SPJ. 2016. Infectious Entry Pathway Mediated by the Human Endogenous Retrovirus K Envelope Protein. J Virol 90:3640–3649.

58. Paterson RG, Russell CJ, Lamb RA. 2000. Fusion protein of the paramyxovirus SV 5: destabilizing and stabilizing mutants of fusion activation. Virology 270:17–30.

59. Hatziioannou T, Valsesia-Wittmann S, Russell SJ, Cosset FL. 1998. Incorporation of fowl plague virus hemagglutinin into murine leukemia virus particles and analysis of the infectivity of the pseudotyped retroviruses. J Virol 72:5313–5317.

60. McKay T, Patel M, Pickles RJ, Johnson LG, Olsen JC. 2006. Influenza M2 envelope protein augments avian influenza hemagglutinin pseudotyping of lentiviral vectors. Gene Ther 13:715–724.

61. Henkel JR, Weisz OA. 1998. Influenza virus M2 protein slows traffic along the secretory pathway. pH perturbation of acidified compartments affects early Golgi transport steps. J Biol Chem 273:6518–6524.

62. Moll M, Klenk H-D, Maisner A. 2002. Importance of the cytoplasmic tails of the measles virus glycoproteins for fusogenic activity and the generation of recombinant measles viruses. J Virol 76:7174–7186.

63. Sandrin V, Boson B, Salmon P, Gay W, Nègre D, Le Grand R, Trono D, Cosset F-L. 2002. Lentiviral vectors pseudotyped with a modified RD114 envelope glycoprotein show increased stability in sera and augmented transduction of primary lymphocytes and CD34+ cells derived from human and nonhuman primates. Blood 100:823–832.

64. Mochizuki H, Schwartz JP, Tanaka K, Brady RO, Reiser J. 1998. High-titer human immunodeficiency virus type 1-based vector systems for gene delivery into nondividing cells. J Virol 72:8873–8883.

65. Simmons G, Wool-Lewis RJ, Baribaud F, Netter RC, Bates P. 2002. Ebola virus glycoproteins induce global surface protein down-modulation and loss of cell adherence. J Virol 76:2518–2528.

66. Salvador B, Sexton NR, Carrion R Jr, Nunneley J, Patterson JL, Steffen I, Lu K, Muench MO, Lembo D, Simmons G. 2013. Filoviruses utilize glycosaminoglycans for their attachment to target cells. J Virol 87:3295–3304.

67. Wrensch F, Karsten CB, Gnirß K, Hoffmann M, Lu K, Takada A, Winkler M, Simmons G, Pöhlmann S. 2015. Interferon-Induced Transmembrane Protein-Mediated Inhibition of Host Cell Entry of Ebolaviruses. J Infect Dis 212 Suppl 2:S210–8.

68. Wickersham IR, Lyon DC, Barnard RJO, Mori T, Finke S, Conzelmann K-K, Young JAT, Callaway EM. 2007. Monosynaptic restriction of transsynaptic tracing from single, genetically targeted neurons. Neuron 53:639–647.

